# Artemisinin-resistant malaria parasites show enhanced transmission to mosquitoes under drug pressure

**DOI:** 10.1101/2020.02.04.933572

**Authors:** Kathrin Witmer, Farah A. Dahalan, Michael J Delves, Sabrina Yahiya, Oliver J. Watson, Ursula Straschil, Darunee Chiwcharoen, Boodtee Sornboon, Sasithon Pukrittayakamee, Richard D. Pearson, Virginia M. Howick, Mara K. N. Lawniczak, Nicholas J. White, Arjen M. Dondorp, Lucy C. Okell, Andrea Ruecker, Kesinee Chotivanich, Jake Baum

**Author notes:** Correspondence to: Jake Baum, Department of Life Sciences, Imperial College London, Exhibition Road, South Kensington, London SW7 2AZ, United Kingdom. contributed equally (Kathrin Witmer, Farah A. Dahalan, Michael J Delves).

## Abstract

Resistance to artemisinin combination therapy (ACT) in the *Plasmodium falciparum* parasite is threatening to reverse recent gains in reducing global deaths from malaria. Whilst resistance manifests as delayed asexual parasite clearance in patients following ACT treatment, the phenotype can only spread geographically via the sexual cycle and subsequent transmission through the mosquito. Artemisinin and its derivatives (such as dihydroartemisinin, DHA) as well as killing the asexual parasite form are known to sterilize male, sexual-stage gametes from activation. Whether resistant parasites overcome this artemisinin-dependent sterilizing effect has not, however, been fully tested. Here, we analysed five *P. falciparum* clinical isolates from the Greater Mekong Subregion, each of which demonstrated delayed clinical clearance and carried known resistance-associated polymorphisms in the *Kelch13* gene (PfK13^var^). As well as demonstrating reduced sensitivity to artemisinin-derivates in *in vitro* asexual growth assays, certain PfK13^var^ isolates also demonstrated a marked reduction in sensitivity to these drugs in an *in vitro* male gamete activation assay compared to a sensitive control. Importantly, the same reduction in sensitivity to DHA was observed when the most resistant isolate was assayed by standard membrane feeding assays using *Anopheles stephensi* mosquitoes. These results indicate that ACT use can favour resistant over sensitive parasite transmission. A selective advantage for resistant parasite transmission could also favour acquisition of further polymorphisms, such as mosquito host-specificity or antimalarial partner–drug resistance in mixed infections. Favoured transmission of resistance under ACT coverage could have profound implications for the spread of multidrug resistant malaria beyond Southeast Asia.

**ONE SENTENCE SUMMARY:** Artemisinin-resistant clinical isolates can also demonstrate resistance to the transmission-blocking effects of artemisinin-based drugs, favouring resistance transmission to the mosquito.

## INTRODUCTION

Malaria kills more than 400,000 people each year (*1*). Whilst there has been a marked reduction in global rates of malaria disease since the new millennium, progress has stalled recently even reversing in some regions (*1*). A critical factor threatening future gains is the emergence and spread of drug resistance in the most virulent parasite *Plasmodium falciparum* (*2*). Of most concern is the reported spread of resistance to frontline artemisinin-based drugs in the Greater Mekong Subregion (GMS) of Southeast Asia (*3, 4*). Artemisinin has revolutionised treatment for severe malaria. The drug acts rapidly to clear the clinical symptoms of malaria by killing the asexual parasite in host red blood cells. Although a precise mechanism of action is contested, it is thought that iron-mediated activation of artemisinin arising from parasite metabolism of haemoglobin causes the drug to be both highly reactive and consumed rapidly in the process of its action(*5*). Consequently, use of artemisinin or its derivatives requires coformulation with longer-lasting partner drugs as artemisinin-based combination therapies (ACTs). In recent years, however, resistance to both artemisinin and partner drugs, including piperaquine and mefloquine, has increased in prevalence throughout Southeast Asia (*4, 6-8*). The spread of such multidrug resistant parasites beyond the GMS region could prove catastrophic for global malaria control.

Resistance to artemisinin is strongly associated with non-synonymous single nucleotide polymorphisms (SNPs) in the propeller domain of *P. falciparum* Kelch 13 (PfK13) (*9*) a protein with multiple likely functions in the parasite cell (*5*). Based on the SNP analysis, several PfK13 variants (PfK13^var^) have been defined displaying different degrees of delayed parasite clearance in patients under ACT treatment. PfK13 variants include mutually exclusive SNPs giving rise to amino acid changes C580Y, R539T, I543T and Y493H (*4, 7, 10, 11*). Whilst the precise mechanism by which PfK13^var^ determines resistance remains ill-defined (*5*), PfK13^var^ parasites show an upregulation in the unfolded protein cell stress response (*12*). Given the importance of this pathway to general cell viability, PfK13^var^ parasites may be better able to deal with stresses arising from drug damage on cell function (*12*). Persistence of parasites in the blood of infected individuals will lead to their delayed clearance and ultimately treatment failure. Among PfK13 polymorphisms, the PfK13^C580Y^ genotype is the most widely spread variant currently circulating in eastern Southeast Asia (*7*).

Drug resistance and its spread is traditionally seen through the prism of disease, in the case of malaria the asexual replicative stages of the life cycle carried in blood circulation. However, resistance can only spread with passage of the parasite through the mosquito, a fundamental step in the *Plasmodium* lifecycle (*13*). Transmission of malaria parasites is solely mediated by non-pathogenic sexual stages called gametocytes. These gametocytes mature over the course of 10-12 days and are the only stages infectious to mosquitoes (*13*). During a mosquito blood feed, male and female gametocytes are taken up and activate in the mosquito midgut into male and female gametes. These activated gametes then fertilise and form a motile zygote (ookinete) that infects the midgut epithelium forming an oocyst on the gut lining (*14*). The oocyst eventually bursts releasing sporozoites that can be transmitted back into humans during a subsequent bite from an infected mosquito.

Whilst the activity of artemisinin derivatives on asexual-stage parasites is well known, one overlooked property of these drugs is their ability to target sexual stages, specifically their ability to block the activation of male gametes (exflagellation), which underpins transmission (*15, 16*). This raises the question as to whether artemisinin-resistant parasites are also resistant to this sterilizing effect in the context of transmission to the mosquito. Here, we sought to test how clinical isolates with demonstrated tolerance or treatment delay against artemisinin (i.e. asexual stage growth) fair in their transmissibility through the mosquito under artemisinin coverage. We show that PfK13^var^ isolates can exhibit transmission resistance, which manifests through an increased ability to activate gametes and infect mosquitoes under artemisinin treatment compared with sensitive controls. These findings have important implications for modelling the spread of resistance across geographical regions. Artemisinin-resistant transmission emphasizes the need for future combination therapies that include a transmission-blocking component if we are to stem the spread of resistance beyond the Greater Mekong Subregion.

## RESULTS

### Selection and adaptation of Southeast Asian *P. falciparum* clinical isolates for *in vitro* study

*P. falciparum* clinical isolates that successfully adapted to long-term culture (Chotivanich, unpublished data) were derived from a previous, multi-centre, open-label, randomised trial collecting samples from patients with acute, uncomplicated malaria (*10*). Among isolates, five were followed further based on their ability to form functional mature gametocytes *in vitro.* These were compared to a standard laboratory control parasite NF54. Each was validated by PCR, confirming the five clinical isolates as having variant polymorphisms in the gene, PfK13^var^ (**Table 1**). Each *P. falciparum* isolate was then tested *ex vivo* for sensitivity to the artemisinin derivative artesunate using the 24-hour trophozoite maturation inhibition assay (TMI)(*17*). While PfK13^var^ isolates presented a wide range of IC_50_ values, they all showed increased resistance to artesunate compared to NF54 and an additional PfK13^WT^ culture-adapted Thai laboratory strain, TM267 (**Table 1**).

**Table 1.** Characteristics of the *P. falciparum* field isolates presented in this study. Clinical isolate codes are shown. IC_50_ values of field isolates assayed with three independent biological replicates using the trophozoite maturation inhibition assay (TMI) (*17*). Standard deviations are indicated in brackets (SD). *P. falciparum* TM267 isolate originate from a previous study (*17*) with others from (*10*).

| Clinical isolate code | Year | Origin | Parasite clearance half-life (hours) | Mean (SD) artesunate IC<sub>50</sub> (ng/ml) | PfK13 genotype (PCR) |
| --- | --- | --- | --- | --- | --- |
| TM267 | 1995 | Thailand | ND | 0.7 (0.5–0.9) | WT (17) |
| ARN1G | May 2011–April 2013 | Ranong, Thailand | 7.1 | 6.8 (0.1) | G449A |
| APS2G | May 2011–April 2013 | Srisaket, Thailand | 6.1 | 9.3 (1.6) | R539T |
| APS3G | May 2011–April 2013 | Srisaket, Thailand | 6.2 | 12.9 (0.4) | R539T |
| APL4G | May 2011–April 2013 | Pailin, West Cambodia | 5.1 | 6.6 (1.0) | C580Y |
| APL5G | May 2011–April 2013 | Pailin, West Cambodia | 7.7 | 5.2 (2.4) | C580Y |

In addition to PfK13 genotype, the genetic background of each parasite isolate was investigated to explore whether additional mutations might be present such as those associated with other drug resistance phenotypes. Whole genome sequencing analysis was completed for each, confirming different PfK13 genotypes (**Table 2**). In addition, multiple previously reported mutations in genes associated with various drug sensitivities were found among PfK13^var^ isolates (as reviewed in(*2*)) (**Table 2**). Mutations were found in the chloroquine resistance transporter (PfCRT) agreeing with reported mutations found in some parasites following ACT treatment(*18*). None of the isolates, however, carried mutations in PfCRT loci recently associated with increased DHA-piperaquine treatment failure (*6, 7, 19*).

**Table 2.**
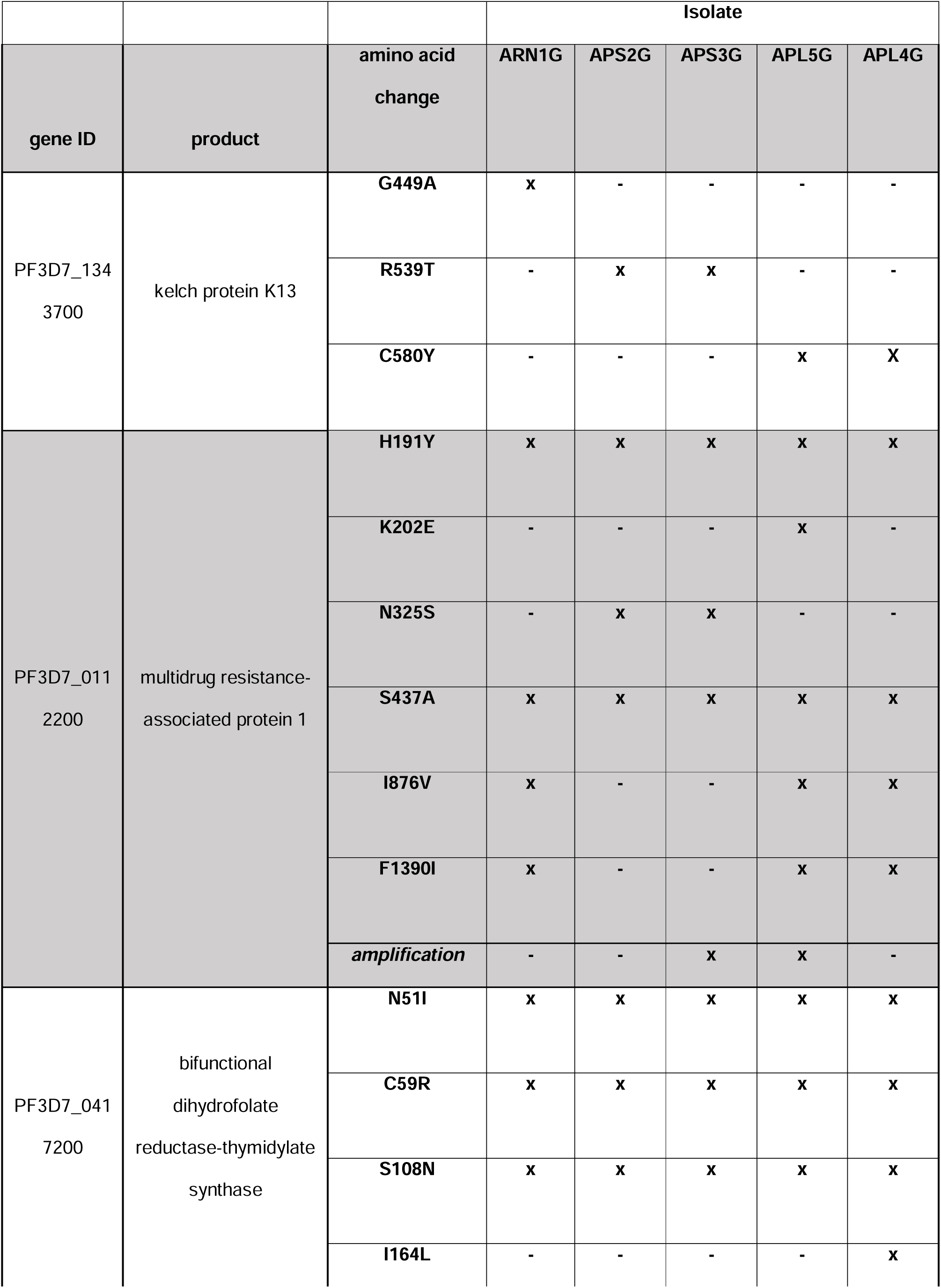
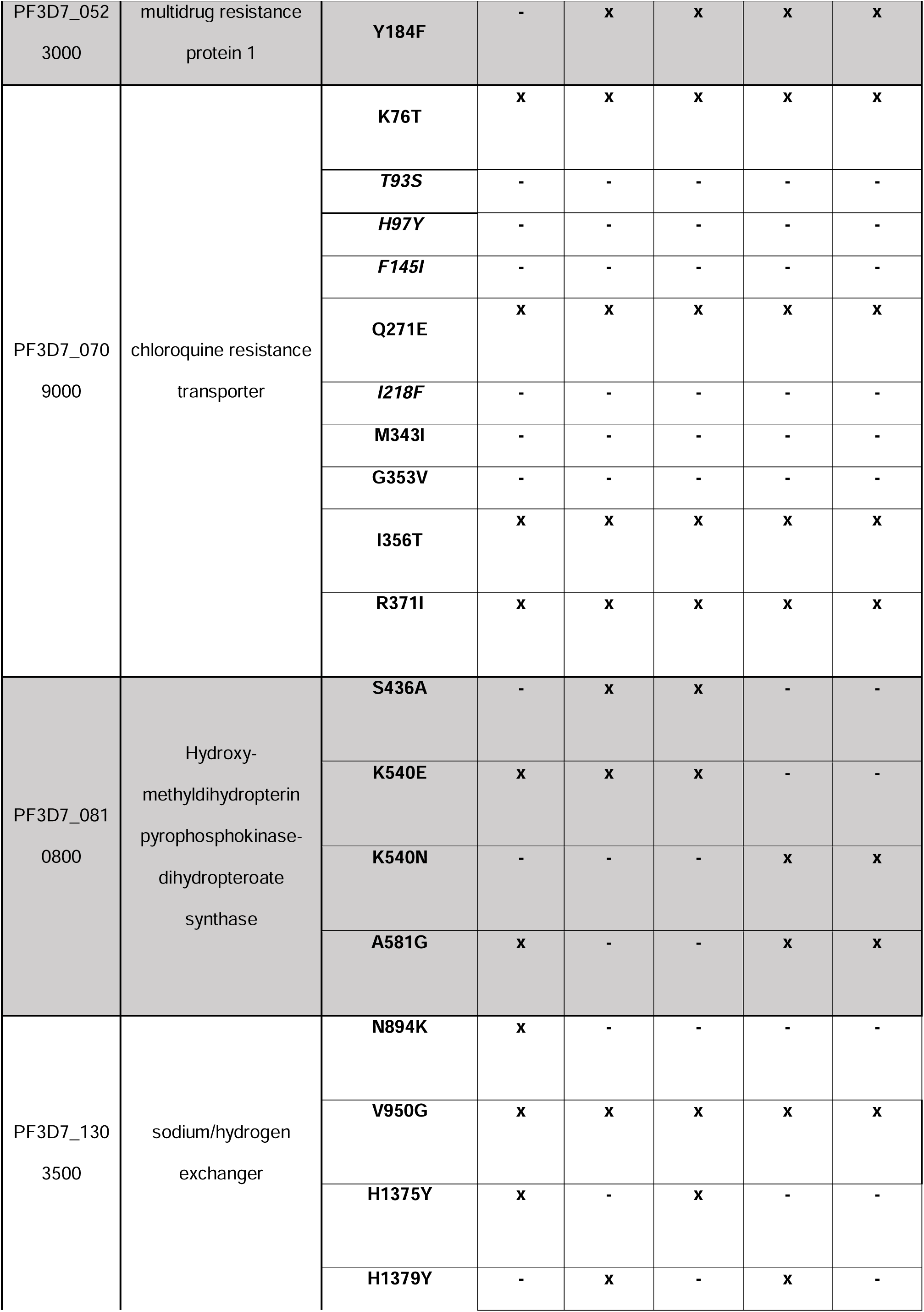
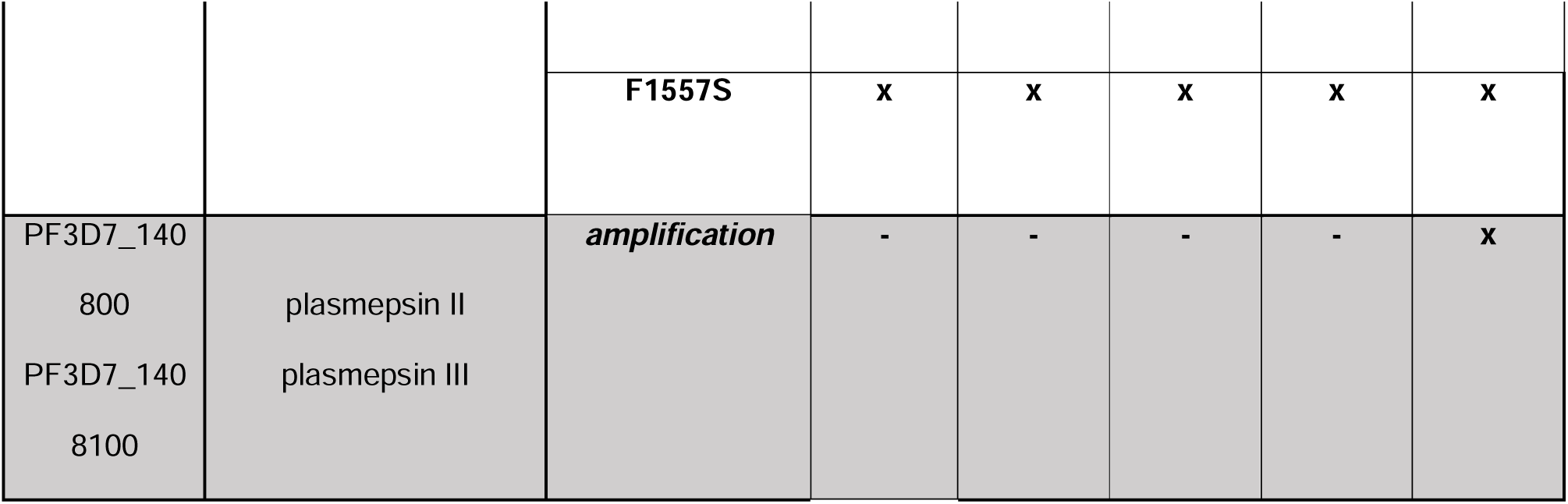
Summary molecular markers associated with antimalarial drug resistance for isolates used in this study. Presence (x) or absence (-) of the polymorphism is indicated for each protein. All amino acid changes in italics have recently been associated with ACT treatment failure within the eastern Greater Mekong subregion.

Increased copy number for the multidrug resistance transporter Pfmdr1 and enzymes plasmepsin II/plasmepsin III are known to be associated with enhanced survival of parasites exposed to mefloquine or piperaquine respectively(*20-23*). To test if any of the field isolates harboured copy number variations, we created sashimi plots of the next generation sequencing coverage (**Figure S1**) and found that plasmepsin II and plasmepsin III are duplicated for isolate APL4G (**Table 2**). APS3G and APL5G both had copy number variants of mdr1 (**Table 2**). This suggests that APS3G and APL5G will likely show resistance to mefloquine, whilst APL4G will likely show resistance to piperaquine (*21, 24*).

### Variation in PfK13 results in a growth defect in asexual blood stages but not in mosquito stages

Polymorphisms in the *PfKelch13* gene (PfK13^var^) that are associated with artemisinin resistance are known to also show reduced asexual blood stage growth(*25, 26*). To validate this in selected isolates, parasites were set up in synchronised ring stage cultures, at a starting parasitaemia of 2%, and followed over the course of eight days. Parasitaemia was analysed every second day by flow cytometry and cultures re-diluted to 2%. NF54 parasites showed a cumulative parasitaemia as expected under standard laboratory conditions (**Figure 1A**). Parasites with a PfK13^var^ showed a significantly reduced replication rate in *in vitro* compared to NF54 (**Figure 1A**) agreeing with previous studies (*25, 26*). To explore the underlying mechanism of slowed growth, we measured the number of merozoites per schizont, which will directly determine potential growth rates(*27*). Late synchronised schizonts were blocked from merozoite egress using the protein-kinase G (PKG) inhibitor, compound 2(*28*). Thin smears, 12 hours later, were then made of each culture and stained with the nuclear stain 4′,6-diamidino-2-phenylindole (DAPI) to count nuclei per schizont. PfK13^var^ isolates displayed fewer nuclei per schizont than the PfK13^WT^ isolate and NF54 control, suggesting that the observed reduced growth rate may at least be partly explained by a reduction in the number of progeny (**Figure 1B**).

**Figure 1.**
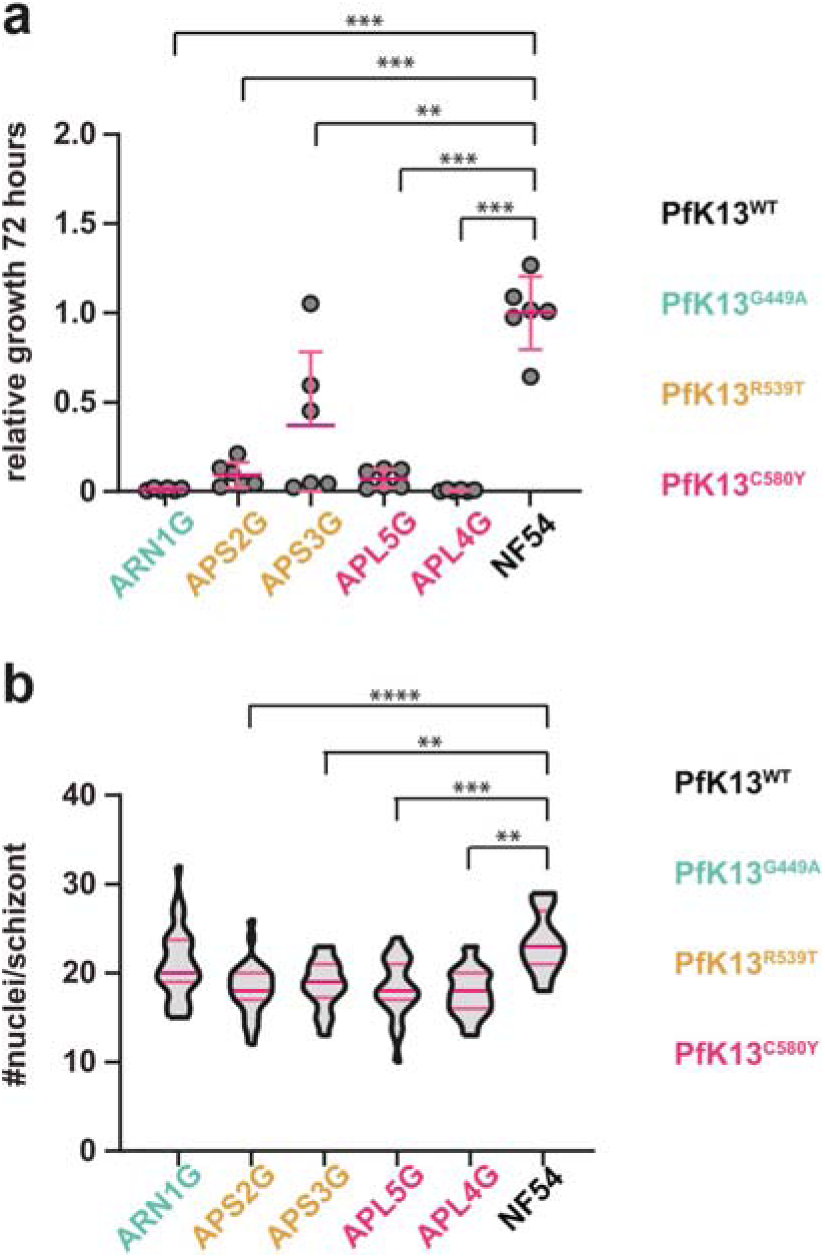
Characterisation of *P. falciparum* clinical isolates. **A.** Relative cumulative growth of *P. falciparum* clinical isolates compared to NF54. Parasitaemia was measured by flow cytometry every other day for eight consecutive days (four replication cycles). Six biological replicates from two parallel experiments are shown. Clinical isolates with K13^var^ grow significantly slower than NF54 (K13^WT^, unpaired t-test, ** p<0.01; ***p<0.0001). **B.** K13^var^ parasites have less nuclei per schizont than K13^WT^ parasites with the exception of isolate ARN1G (unpaired t-test ** p<0.01; ***p<0.0001).

To investigate the transmission capability of each *P. falciparum* field isolate, we induced gametocytes at a starting parasitaemia of 2% (*29*) and, 14 days post induction, fed cultures to *Anopheles stephensi* mosquitoes by standard membrane feeding assay (SMFA)(*30*). No significant differences were noted in the stage V (mature) gametocytaemia for isolates (in terms of relative numbers of gametocytes to asexual parasites). However, upon activation, male exflagellation rates were reduced in PfK13^var^ isolates compared with PfK13^WT^ (**Figure S2**). Ten days post-feeding, mosquito midguts were dissected, and oocysts numbers recorded. All field isolates were found to be capable of infecting mosquito midguts at varied intensity levels, i.e. oocyst counts per midgut, as reported previously for Cambodian field isolates (*31*). To test if PfK13^var^ led to a reduced replication rate in mosquito stage growth (following the reduced merozoite count), we measured the diameter of each oocyst in these infections as a proxy for replication. Oocyst size showed no consistent pattern of variation when compared to controls other than a PfK13^R539T^ variant which displayed significantly larger oocysts than the PfK13^WT^ control (**Figure S2**). This shows that whilst variations in PfK13 may reduce parasite multiplication rate in the asexual blood stage it does not appear to directly influence transmission and growth in the mosquito-stage.

### Exflagellation sensitivity to different antimalarials among isolates with varied PfK13 genotypes

It has previously been shown that artemisinin and its derivatives have an inhibitory effect on male gamete exflagellation, irreversibly sterilizing male gametocytes from activation (*15, 16*). To explore whether PfK13^var^ isolates were resistant to this sterilizing effect we tested mature gametocyte culture capacity to activate in the dual gamete formation assay (PfDGFA) (*15, 16*). 24-hour incubation of cultures with the artemisinin derivative dihydroartemisinin (DHA) was found to be insufficient to elicit a complete inhibition of exflagellation for PfK13^WT^ NF54 parasites (**Figure S3**), likely as a result of the rapid instability of the drug (*32*). To improve activity and allow for a comparative analysis between isolates, gametocytes were exposed to a second compound dose 24 hours after the first, resulting in a double-dose regimen with a readout after 48 hours. Double exposure consistently gave complete inhibition of male activation with DHA at the highest concentration tested (**Figure S3**). Female gamete activation was unaffected as found previously (*15*). In parallel, two other artemisinin derivatives and four other antimalarial drugs were tested by PfDGFA (**Figure S4** and **Figure S5**). Exflagellation rates for PfK13^var^ isolates varied in the presence of drug, with two PfK13^R539T^ and PfK13^C580Y^ isolates consistently showing tolerance to artemisinin-derivatives (**Figure 2A**). The lack of a consistent pattern of reduced sensitivity to artemisinin-based drugs across PfK13^var^ isolates suggests that K13 polymorphisms alone likely do not completely explain sensitivity of sexual stages to artemisinin treatment. This observation is corroborated by similar findings with piperaquine resistance, which is mostly, but not always, explained by copy number variants in plasmepsin II and plasmepsin III (*21*). However, the presence of even a single isolate with both reduced sensitivity to artemisinin-based drugs in asexual and sexual stages, suggests that there is the potential that a resistant strain at treatment might be favoured for transmission to the mosquito.

**Figure 2.**
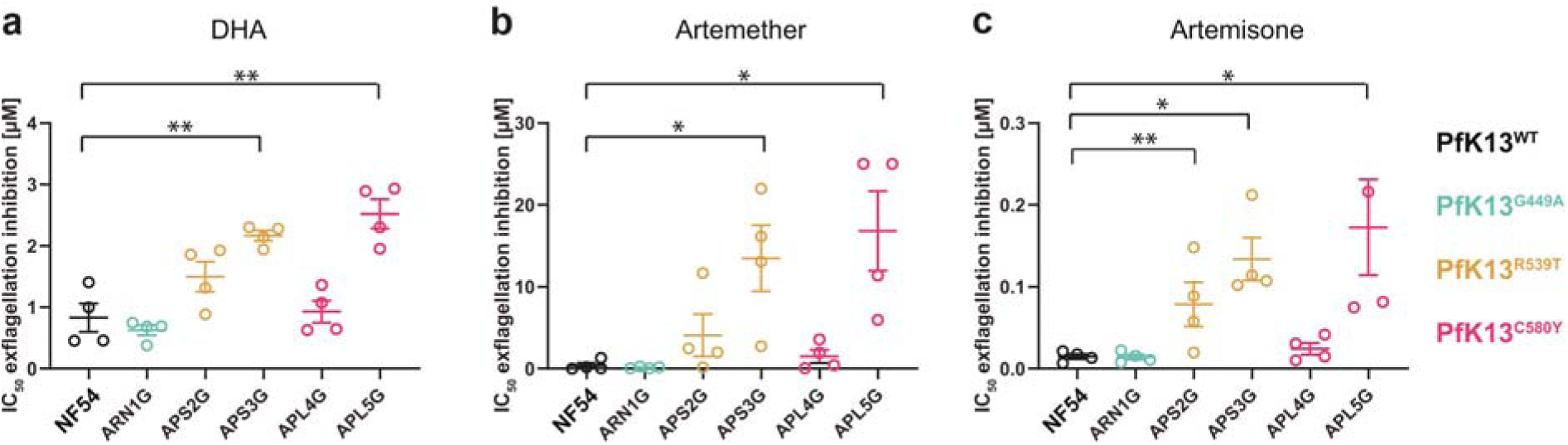
Exflagellation inhibition reported as IC_50_ values of clinical isolates. Two parasite isolates (APS3G and APL5G) show consistent resistance to sterilizing effects of three different artemisinin-related drugs, dihydroartemisinin (DHA) (**A**), Artemether (**B**) and Artemisone (**C**) on exflagellation compared to the NF54 PfK13^WT^ control. One additional isolate (APLS2G) shows increased resistance to Artemisone. IC_50_ values were compared to NF54 (unpaired t-test * p<0.05; ** p<0.01)

### Transmission of field isolates with different PfK13 genotypes under DHA drug selection

To explore the hypothesis that PfK13^var^-associated resistance might allow resistant parasites to more efficiently infect mosquitoes under drug coverage, we selected the PfK13^C580Y^ isolate APL5G. This isolate showed a comparable level of mosquito infection to NF54 (**Figure S2**) and also showed a high level of male gamete activation resistance to DHA (**Figure 2**). Gametocyte cultures of both parasites were exposed to different concentrations of DHA for 48 hours using our double-dosing regimen, before feeding to *An. stephensi* mosquitoes by SMFA. At day 10 post-feed, mosquitos were dissected, and midguts examined for oocyst load (**Figure S6**). Generalised linear mixed effects models were used to analyse infection intensity (number of oocysts per midgut) and infection prevalence (proportion of midguts with oocysts) in response to treatment with DHA, in order to incorporate data from 18 individual SMFA experiments within the same modelling framework. A decrease in both intensity and prevalence of infected mosquitoes was observed for both parasite isolates with increasing DHA concentration (**Figure 3A** and **Figure 3B**). A significant decrease in both the oocyst intensity (ratio of oocyst intensity = 0.73, 95% CI: 0.66-0.80) and prevalence (odds ratio = 0.46, 95% CI: 0.41-0.52) of mosquito infection was observed for NF54 with increasing drug concentration. In contrast, APL5G parasites (C580Y) showed no evidence for a significant decrease in oocyst intensity with increasing DHA (ratio of oocyst intensity = 0.84, 95% CI: 0.65-1.07). Increasing concentrations of DHA did still reduce the oocyst prevalence for APL5G (odds ratio = 0.72, 95% CI: 0.56-0.96), however, this effect was significantly less than the effect seen for WT parasites (**Table 3**).

**Table 3.** Effects of DHA treatment on oocysts prevalence and intensity. Analysis was done using generalised linear mixed effects models to incorporate 18 SMFA experiments into a single analysis.

|  | Prevalence of Infection |  |  | Oocyst Intensity* |  |  |
| --- | --- | --- | --- | --- | --- | --- |
|  | *Odds Ratio* | *95% CI* | *p* | *Ratio of oocyst intensity* | *95% CI* | *p* |
| K13<sup>var</sup> in the absence of DHA | 0.20 | 0.16 – 0.25 | <0.001 | 0.17 | 0.13 – 0.23 | <0.001 |
| DHA (µM): WT line | 0.46 | 0.41 – 0.52 | <0.001 | 0.73 | 0.66 – 0.80 | <0.001 |
| DHA (µM): K13<sup>var</sup> | 0.72 | 0.56 – 0.96 | <0.001 | 0.84 | 0.65 – 1.07 | 0.069 |

\* Negative binomial overdispersion parameter: 0.837

**Figure 3.**
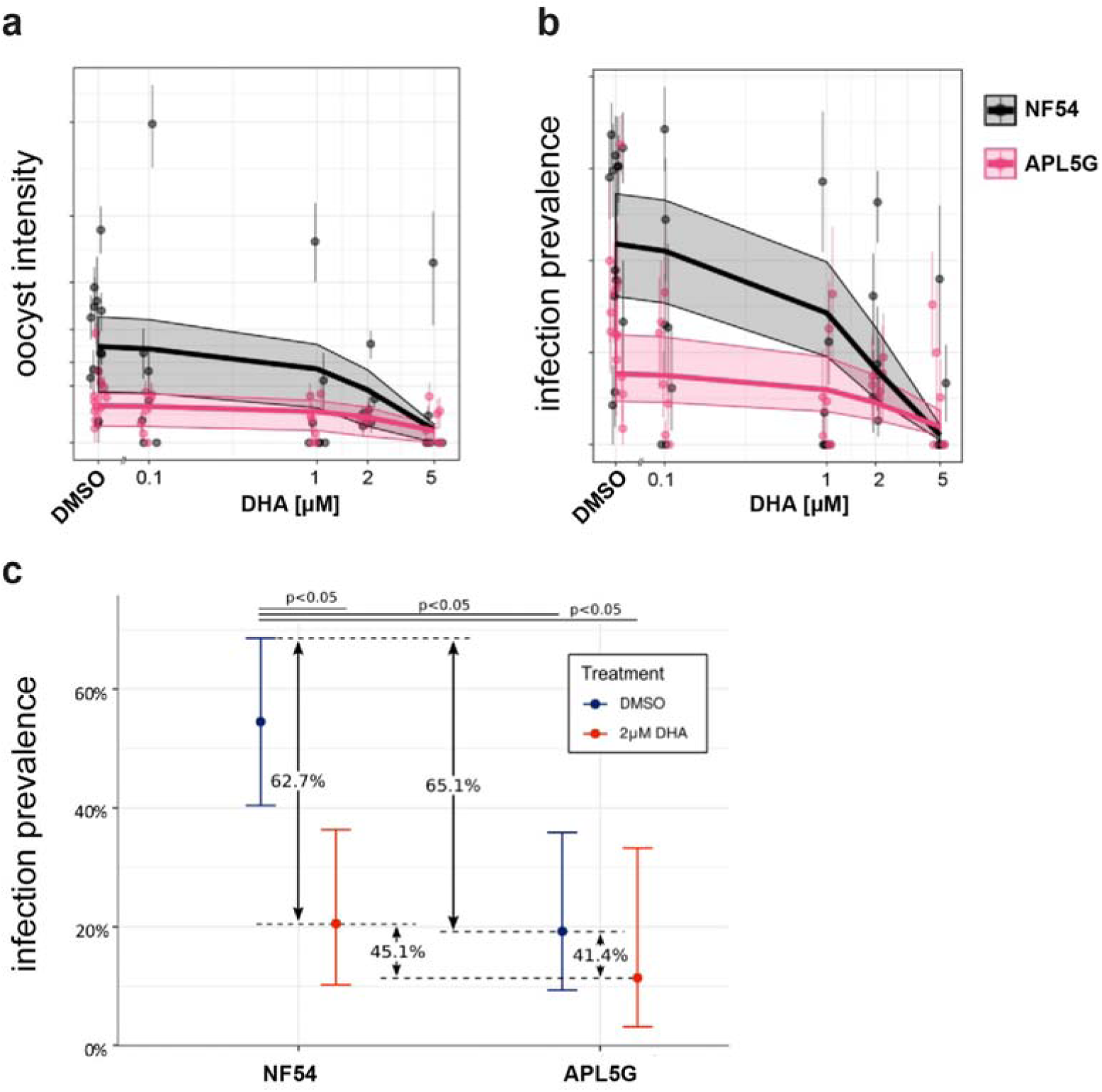
The impact of DHA on the PfK13^WT^ and PfK13^C580Y^ transmission potential. Graphs show the overall results of oocysts count from 18 individual SMFA experiments oocysts infection intensity (**A**) and infection prevalence (**B**) of the sensitive isolate PfK13^WT^ (NF54) versus the artemisinin resistant PfK13^C580Y^ isolate (APL5G) after incubation with either DMSO (no DHA) and DHA at specified concentrations. Points and whiskers on each plot show mean and bootstrapped 95% CI for each replicate, with the predicted relationship and 95% CI shown with the trend line and shaded region. In the absence of DHA (DMSO), APL5G is predicted to produce significantly fewer oocysts and infections, whereas in the presence of DHA concentrations greater than 2 µM DHA, the transmission potential of PfK13^C580Y^ is comparable to NF54/PfK13^WT^. **C**. Fitness costs associated with DHA resistance. The relative reduction in infection prevalence due to DHA treatment in NF54 is greater (62.7%) than APL5G (41.4%), which suggests that APL5G is significantly more likely to infect mosquitoes under drug treatment (p<0.05) compared to the absence of drug.

To position these findings in the context of likely transmission events, we further explored the impact of DHA on transmission at a single concentration (2µM) of DHA in comparison with DMSO. 2µM reflects the likely peak of DHA serum concentration when following the recommended WHO dose for the ACT, DHA-piperaquine (*33-35*). In the absence of DHA, NF54 parasites were consistently observed to have an increased infection prevalence (55.1%, 95% CI: 50.5% - 57.8%) compared to APL5G parasites (19.2%, 95% CI: 13.3% - 26.0%) i.e. all things considered, NF54 transmits better in the absence of drug (**Figure 3C**). This suggests that a fitness cost is associated with the APL5G genotype, which causes a sizeable reduction in the onward probability of infection relative to WT parasites in the absence of DHA. However, this changes sigifnificantly in the presence of drug. With 2µM DHA, no significant difference was observed between NF54 parasites (20.6%, 95% CI: 27.7% - 14.3%) and APL5G parasites (12.1%, 95% CI: 4.1% - 25.4%) (**Figure 3C**) demonstrating a profound impact on NF54 but not APL5G parasites. The decreased impact of DHA on the ability of the K13^var^ isolate to infect mosquitoes indicates that the resistance polymorphism significantly increases the transmission potential of the parasite in the presence of DHA. This observed transmission resistance phenotype may offset any fitness costs (such as growth) observed in the absence of drug.

## DISCUSSION

The threat of spreading artemisinin resistance to the treatment of malaria disease has focussed global attention on the mechanisms underlying resistance in the parasite, *Plasmodium falciparum*. However, only limited focus has been placed on how resistant parasites transmit through the *Anopheles* mosquito vector. In this paper, we have shown clear evidence that certain clinical isolates, with defined artemisinin resistance based on the known PfKelch13 marker (in terms of clinical delayed clearance and reduced asexual growth sensitivity in the presence of artemisinin-based drugs) are also able to transmit better to mosquitoes under drug coverage compared to artemisinin-sensitive controls. The molecular basis of this transmission-resistance phenotype is likely complex and is clearly not defined simplistically by PfKelch13 alone. Ongoing studies using CRISP/Cas9 gene editing(*26*) may be able to address the polygenic nature of this phenotype. However, the disconnect between PfK13 and transmission-resistance is clear from the observation that of the five clinically resistant isolates tested, whilst each showed clear resistance to artemisinin-based drugs in asexual growth, there was varied sensitivity in transmission stages. However, the transmission-resistance phenotype is nonetheless robust for certain isolates.

As with previous studies, we first started with investigation of the asexual growth rate, confirming a consistent reduced rate in resistant isolates. The asexual growth rate reduction seen in PfK13^var^ isolates likely acts as both a selective cost for parasite growth (in being out competed in normal infections) but also likely explains how these parasites persist during drug treatment, i.e. explaining delayed clearance(*5*). Switching our focus to sexual commitment and development, we next explored gametocyte production. With the caveat that different parasite isolates always show marked differences in gametocyte formation capacity, we did not observe any obvious reduced capacity among PfK13^var^ isolates in gametocyte production. All five field isolates produced equivalent numbers of mature gametocytes (stage V gametocytes) after day 14 upon induction. Indeed, the reverse correlation between drug resistance and sexual commitment has been consistently reported. Clinical isolates with demonstrated drug resistance and delayed clearance have been consistently reported to produce a higher gametocytaemia, suggesting a potentially elevated potential for transmission(*10*).

Having confirmed capacity to form gametocytes, we assessed the capacity to generate exflagellation centres, a marker of male gamete activation capacity in the mosquito. We found no direct correlation between gametocytaemia and exflagellation count. This lack of correlation may be due to differences in the sex ratio between parasite isolates (reduced males mean less exflagellation centres though gametocytaemia may be the same). Whilst commitment of gametocytes to either male or female is poorly understood, it is entirely conceivable that gametocytes mature or differentiate into either male or female at differing rates in each isolate. Unfortunately, sex ratios were untested here due to a paucity of markers for male and female gametocytes and challenges with definitive differentiation of the sexes using Giemsa stain. Irrespective, gametocyte conversion rates have been shown to be sensitive to asexual stage replication, which itself is affected by drugs. This suggests that there is the potential for a trade-off between asexual stages and sexual stages in ensuring the spread of the artemisinin resistant parasites(*36*).

With sexual commitment and exflagellation *in vitro* seemingly uncompromised in resistant isolates we next sought to explore transmissibility directly. We saw no obvious defect in either the transmission capacity (number of oocysts) or the transmission replication rate (as measured by oocyst size) among the five parasite field isolates and NF54. This latter point is noteworthy since it is clear that artemisinin-resistant parasite isolates show a lower asexual growth rate and merozoite (progeny) rate (**Figure 1**), however, number of oocysts and sporogony in the mosquito doesn’t appear to be affected. Thus, PfK13^var^ parasites appear able to commit to sexual reproduction, activate and transmit to mosquitoes at levels that don’t differ dramatically to those commonly seen in sensitive parasites (i.e. beyond variability usually seen between isolates).

Shifting our attention to transmission under drug coverage, tests of the viability of gametocytes for gamete activation using the dual gamete formation assay (PfDGFA) with artemisinin derivatives; DHA, Artemether and Artemisone clearly found that certain PfK13^C580Y^ and PfK13^R539T^ parasites demonstrated significantly higher resistance compared to sensitive controls. Of note, whilst undertaking this work, a parallel study made similar observations. Testing male exflagellation sensitivity to DHA in unrelated culture-adapted PfK13^var^ Cambodian field isolates, Lozano et al found that PfK13^var^ isolates showed a reduced sensitivity of exflagellation rates to DHA treatment, though onward mosquito infectivity was not tested(*37*). Extending this observation to transmission directly we took the most competent transmissible field isolate representing the resistant phenotypes (APL5G, C580Y) compared to the laboratory reference strain NF54, and tested whether transmission resistance plays out in terms of capacity to infect mosquitoes over a range of drug concentrations. Controlling for gametocytaemia, number of cells and haematocrit level for each, infection prevalence in mosquitoes could then be tested and compared between lines. Of note, the infection intensity (number of oocysts found in each mosquito) was consistently different between NF54 and APL5G, as it is for each different culture-adapted parasite strain (see (*31*)). These differences make direct measures of mixed infections challenging. Nonetheless, we found that artemisinin resistant, PfK13^C580Y^ (APL5G) was consistently more likely to transmit malaria under drug pressure compared to its DMSO treated controls than the control NF54 parasite (**Figure 3C**). This was due to the greater impact of DHA on oocyst infection exhibited by the wild-type isolate, which served to offset the decreased transmission potential for APL5G in the absence of artemisinin. This demonstrates that the artemisinin resistant phenotype of APL5G impacts both on asexual blood stages and during transmission to the mosquito in the presence of artemisinin drug.

Although the numbers are small here, the implications are that in the context of a mixed infection, a resistant parasite may be more likely to survive ACT treatment and its gametocytes may be more likely to transmit to the mosquito. Thus, ACT coverage in the field may be favouring, even driving, artemisinin-resistant parasite persistence and transmission. This could explain an important part of the selection of PfKelch mutants observed in the field. For instance, the F446I PfKelch mutation results in only a slight prolongation in parasite clearance half-life and is not associated with ACT treatment failure (*10*). Yet, there is clear selection of this genotype in Myanmar, which could be explained by preferential transmission under artemisinin drug pressure. The effect on outcrossing is also worth considering. Because the sterilizing effects of artemisinin-based drugs appears to be biased towards the male gametocyte (*15, 16*), there is the very real potential that ACT usage in the context of a mixed infection might favour acquisition of other selectively advantageous mutations during transmission. Since the female gametocytes remain unaffected, successful transmission under ACT coverage would likely favour either resistant parasite selfing or mating between resistant males and sensitive females. It is clear from our own usage of a central Asian mosquito vector (*An. stephensi*) and the work of others using the major African vector, *An. coluzzii* (*31*) that artemisinin-resistant parasites can infect non-native mosquitoes. Thus, in a mixed infection where local parasites show a degree of geographical vector adaptation (*38*) an invasive resistant parasite, otherwise at a disadvantage (reduced vector adaptation and slower asexual growth), may acquire a key advantage under ACT coverage in terms of its ability to both transmit and acquire necessary adaptive mutations via recombination with sensitive females. Importantly, this may play out even without a decline in cure rates if transmissibility of the treated infection is increased, such as in high intensity transmission areas at the early stages of resistance invasion before partner drug resistance has emerged. Mixed infection studies *in vivo* and modelling of drug coverage effects with different rates of transmission intensity are clearly needed to explore the implications of transmission resistance in various invasive settings.

Ultimately, these data stress the importance of considering transmission in the context of drug resistance spread and argue strongly for the inclusion of a parasite transmission-blocking component in future antimalarial combination therapies or control strategies.

## Supporting information

Supplementary Data Sheet 1 (DGFA)

Supplementary Data Sheet 2 (SMFA)

## Author contributions

MJD, AR, KC and JB conceptualised the study; KW, FAD, MJD and AR designed experiments; experiments were undertaken by KW, FAD, MJD, SY, US, SC, and BS; RDP, VMH, MKNL, KW and AR generated and curated the genome data; modelling components were designed and executed by OJW and LO. SP, NJW, AD, KC supervised collection of clinical isolates used in the study. KW, FD and JB wrote the manuscript. All authors contributed to overall editing and manuscript approval.

## Data availability

Raw experimental data are available on request, genome data is publicly available at the European Nucleotide Archive (https://www.ebi.ac.uk/ena).

## Supplementary Materials for

### MATERIAL AND METHODS

#### *P. falciparum* asexual blood stage and gametocyte maintenance

Asexual blood stage and gametocytes were cultured as previously described (*29*) with the following modifications: Asexual blood stage cultures were maintained in asexual culture medium (RPMI 1640 with 25 mM HEPES (Life Technologies), 50 µg L^-1^ hypoxanthine (Sigma), 5% A+ human serum (Interstate Blood-Bank) and 5% AlbuMAX II (Life Technologies)). Gametocyte cultures were maintained in gametocyte culture medium (RPMI 1640 with 25 mM HEPES (Life Technologies), 50 µg L^-1^ hypoxanthine (Sigma), 2 g L^-1^ sodium bicarbonate (Sigma), 5% A+ human serum (Interstate Blood-Bank) and 5% AlbuMAX II (Life Technologies)).

#### Mosquito rearing

*Anopheles stephensi* mosquitoes were reared under standard conditions (26°C-28°C, 65%-80% relative humidity, 12 hour:12 hour light/darkness photoperiod). Adults were maintained on 10% fructose.

#### Whole genome sequencing

Genomic DNA isolate and whole genome sequencing, calling of single nucleotide variants and gene amplification copy number variants were undertaken essentially as recently described(*39*).

#### Flow cytometry

For the growth assay, asexual parasites were sorbitol-synchronised at least twice 16 hours apart to create an 8-hour growth window. Starting parasitaemia was seeded at 1-2% early ring stages in triplicates that were treated separately. The assay was performed twice using 3 replicates each. Every other day, parasites were fixed in 4% formaldehyde and 0.2% Glutaraldehyde for at least 10 minutes. After washing with PBS, and DNA was stained with SybrGreen1 (diluted 1:10’000) in the dark for 20 minutes at room temperature. After incubation, cells were washed three times with PBS and resuspended in 80ul PBS. Flow cytometry was performed counting a total of 100’000 cells per condition.

#### Nuclei count

Parasites were synchronised twice using 5% Sorbitol to obtain a 10-hour life cycle window. 10 µM compound 2 was added to late trophozoite stages for a maximum of 12 hours to block egress of the red blood cells (RBC) (*28*). Resulting segmented schizonts were thinly smeared, fixed with 4% Formaldehyde and 0.2% Glutaraldehyde for 20 minutes. Smears were then stained with 1 µg/ml DAPI for 5 minutes and mounted in Vectashield (Vector Laboratories). Z-stacks were taken using a Leica microscope at 100x magnification. Nuclei of arrested segmented schizonts were counted using the plugin tool “Manual counting” on ICY (*40*). Only singly-invaded RBC were counted.

#### Trophozoite maturation assay (TMI)

The trophozoite maturation assay was performed according to (*17*). Briefly, *P. falciparum* infected blood was collected into heparin tubes and centrifuged at 800 g at 4°C for 5 minutes to allow the removal of the plasma and buffy coat. This was followed by three washes in RPMI 1640 (without serum supplement) and adjusted to 3% cell suspension in 10% A+ human serum supplemented RPMI 1640. 96-well microtiter plates (Nunc™ MicroWell™ 96-Well Microplate; Thermofisher Scientific) were predosed with Artesunate dissolved in 5% NaHCO_3_ (Guilin Pharmaceutical Co., Ltd., China), ranging from 0.01 to 400 ng/ml final concentration or no drug as negative control. A 75 µl *P. falciparum* ring stage infected RBC cell suspension was added to the test plate and incubated for 24 hour at 37°C in 5% CO_2_. All samples were tested in triplicate. Upon completion of drug exposure thick and thin blood smears were prepared of all wells and the number of 24 to 30 hour trophozoites (*41*) was counted per 100 infected red blood cells. To identify the inhibition activity of artesunate the percentage of trophozoite maturation compared to the negative control was assessed. The IC_50_ (50% inhibitory concentration) was calculated as the drug concentration causing 50% inhibition of *P. falciparum* maturation from ring stage to trophozoite stage and normalised to the negative control wells. All IC_50_s were determined by sigmoid curve fitting using WinNonlin computer software (version 3.1; Pharsight Corporation, USA). As technical control all *ex vivo* assays were performed in parallel to the standard laboratory Thai strain, TM267.

#### Dual gamete formation assay (DGFA) double-dose format

The double dose DGFA was adapted from previously described methods (*16*) to incorporate an additional drug dosage, accounting for the low compound half-lives of artemisinin and its derivatives. Briefly, compounds were prepared in 10 mM DMSO stocks, and dispensed in serial dilutions into multiwell plates using a HP D300 Digital Dispenser. Samples were normalised to 0.25% DMSO and contained 0.25% DMSO and 12.5 µM Gentian Violet as negative and positives controls, respectively. Half the maximal DMSO content was plated per plate, accounting for the accumulation of DMSO over two dosages. Mature gametocytes with an exflagellation rate of >0.2% of total cells were diluted in gametocyte culture medium to 25 million RBCs per mL. Mature gametocyte culture was plated in drugged 96-well plates and incubated in a humidified chamber under 92% N_2_/5% CO_2_/3% O_2_ (BOC special gases) at 37°C for 24 h. For the second drug dosage at 24 h, the drugged culture was transferred to a second drugged well plate and incubated for a further 24 h under 92% N_2_/5% CO_2_/3% O_2_ at 37°C in a humidified chamber.

At 48 h, gametogenesis was induced with ookinete medium (RPMI 1640 with 25 mM HEPES (Life Technologies), 50 µg L^-1^ hypoxanthine (Sigma), 2 g L^-1^ sodium bicarbonate (Sigma) and 100 µM xanthurenic acid) prepared with 0.5 µg ml^-1^ anti-Pfs25 clone 4B7 (BEI Resources) conjugated to Cy3 (GE Healthcare). Plates were immediately incubated at 4 °C for 4 min and then 28 °C for 5 min before transferring to a Nikon Ti-E widefield microscope. Exflagellation events were recorded by automated phase contrast microscopy, in either 384-well or 96-well plates. For 384-well plate assays, 10-frame time lapses (4 frames sec^-1^) were recorded at x4 objective and1.5x zoom, and for 96-well plate assays, 20-frame time lapses were recorded at x10 magnification and 1.5x zoom. Plates were then protected from light and incubated at 28 °C for 24 h before the automated capture of female gamete formation at the same magnification, utilising single frame fluorescence microscopy. Exflagellation events and female gamete counts per well were derived using an automated ICY Bioimage Analysis algorithm. Resulting counts were converted to percentage inhibition values, calculated relative to positive (C1) and negative (C2) controls:

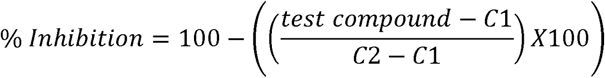

Raw data demonstrated a Z’ factor ≥ 0.4 and was derived from n ≥ 2 and n ≥ 3 technical and biological replicates, respectively. GraphPad Prism (version 8) was used to calculate IC_50_s from the dose response data using with the log(inhibitor) vs. response – variable slope (four parameters) function. IC_50_s were derived from curves demonstrating R^2^ ≥ 0.95.

#### Standard membrane feeding assay

Gametocytes were induced and maintained as mentioned above. At day 14 post induction, gametocytes were spun down at 38°C and resuspended in 5 ml of suspended animation buffer (SA) (*42*). To ensure that a consistent number of RBCs are used for drug incubation, gametocytes were MACS-purified and resuspended in gametocyte medium with 25E6 fresh RBCs. DHA was added to desired end concentration into a 10 ml gametocyte culture and added again 24 hours later (double-dosing within 48 hours). After 48 hours, the parasite culture was mixed with fresh blood and human serum and fed to adult *An. stephensi* mosquitoes using a 3D printed feeder(*30*).

#### Oocyst counts and size

At day 10 post feeding, mosquitoes were dissected, and midguts were stained in 0.1% Mercurochrome and inspected using light microscopy with 10x magnification to count oocysts. To measure oocysts size, midguts of *An. stephensi* fed on *P. falciparum-*infected blood were dissected and fixed with 4% formaldehyde, permeabilised with 0.1% Triton X-100 for one hour, blocked with 3% BSA for 30 minutes and stained with 1 µg/ml in DAPI for 3 minutes. Midguts were washed with 1xPBS and mounted in Vectashield. Images were acquired on a Nikon Ti-Eclipse inverted fluorescence microscope. Images of *P. falciparum*-infected midguts were captured using the DAPI channel and Z-stack imaging to obtain greater depth of oocysts. These stacked images were then processed in ND Processing using the Maximum Intensity Projection option which then create an image with brighter intensity of the oocysts in every midgut. Oocysts detection was automated by using the Automated Spot Detection programme based on the intensity of the oocysts compared to midgut cells (NIS-Elements). The size, diameter and intensity of each selected oocysts were generated in an excel file for analysis.

#### Statistical Modelling of Oocyst infection Intensity and Prevalence

To assess the impact of artemisinin on the ability of each parasite line to form oocysts, we used generalised linear mixed effects models in order to incorporate data from different experimental replicates within the same modelling framework. These models have previously been used to model transmission blocking interventions (*43*). We modelled either oocyst intensity or prevalence as the response with treatment (DHA concentration) included as a fixed effect and 0 µM DHA represented by control groups treated with DMSO. The parasite line treated (PfK13^WT^ or PfK13^C580Y^) was included as a fixed effect to assess the differential impact of artemisinin on transmission success. The impact of treatment between experimental replicates was allowed to vary at random between replicates. A logistic regression (binomial error structure) was used to model the prevalence of mosquito infection, i.e. the presence or absence of oocysts, and a zero-inflated negative binomial distribution was used to model the intensity of infections, i.e. the numbers of mosquito oocysts (*44*). 95% confidence interval estimates were generated for the impact of drug concentration by bootstrapping methodology (with 100,000 replicates).

## Supplementary Figures

**Figure S1.**
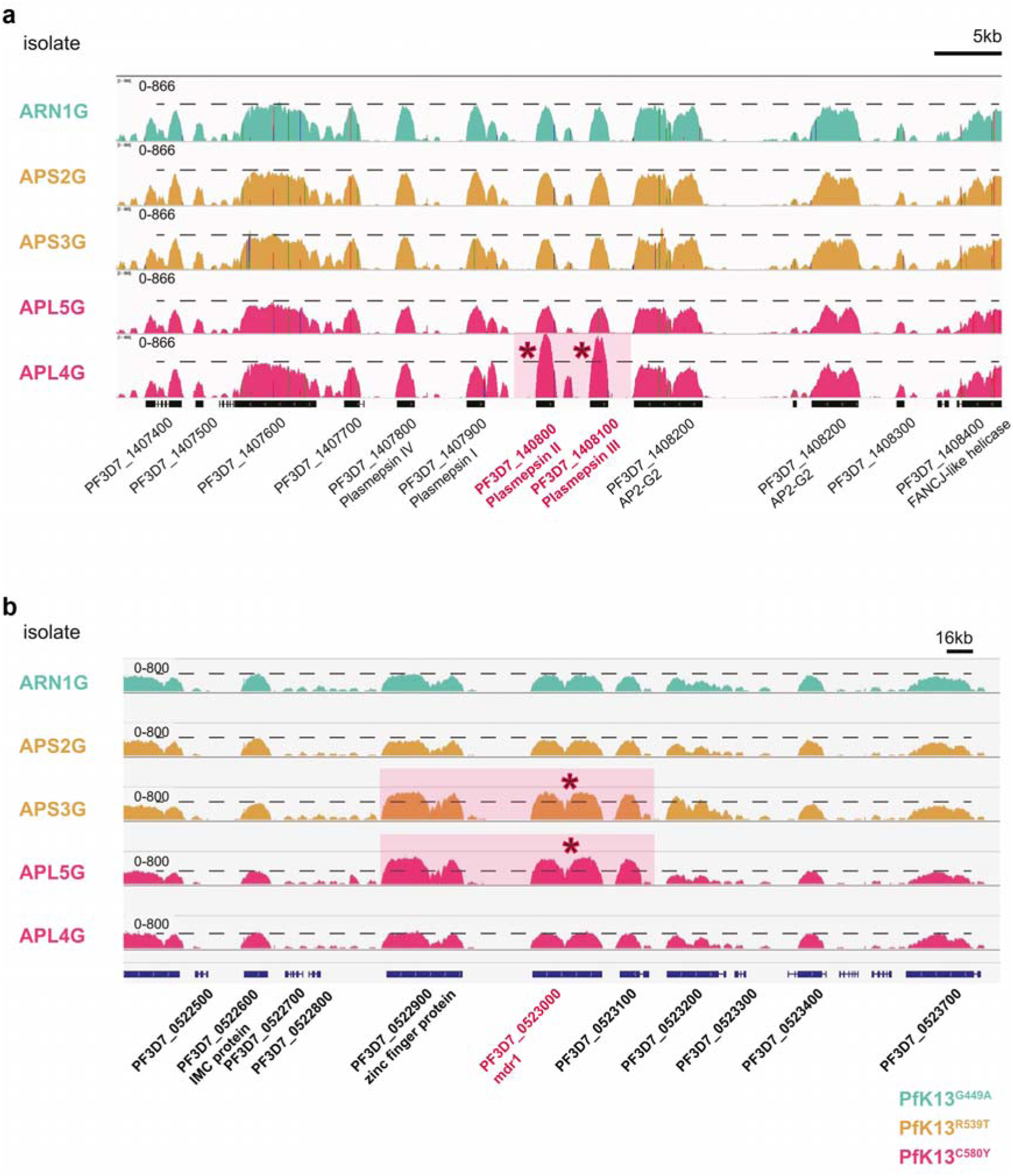
Sashimi plots of field isolates displaying DNA-sequencing coverage over a genomic region of chromosome 5 (**A**) and chromosome 14 (**B**). **A**. *Mdr1* is highlighted in pink, and asterisks indicate a genome duplication event for the gene in isolates APS3G and APL5G. **B**. Plasmepsin II and plasmepsin III are highlighted in pink, and asterisks indicate a genome duplication event for these two genes in isolate APL4G. Dashed line indicates raw DNA-seq pileup reads for the surrounding genes. Parasite isolates and sashimi plots are highlighted according to PfK13 variant.

**Figure S2.**
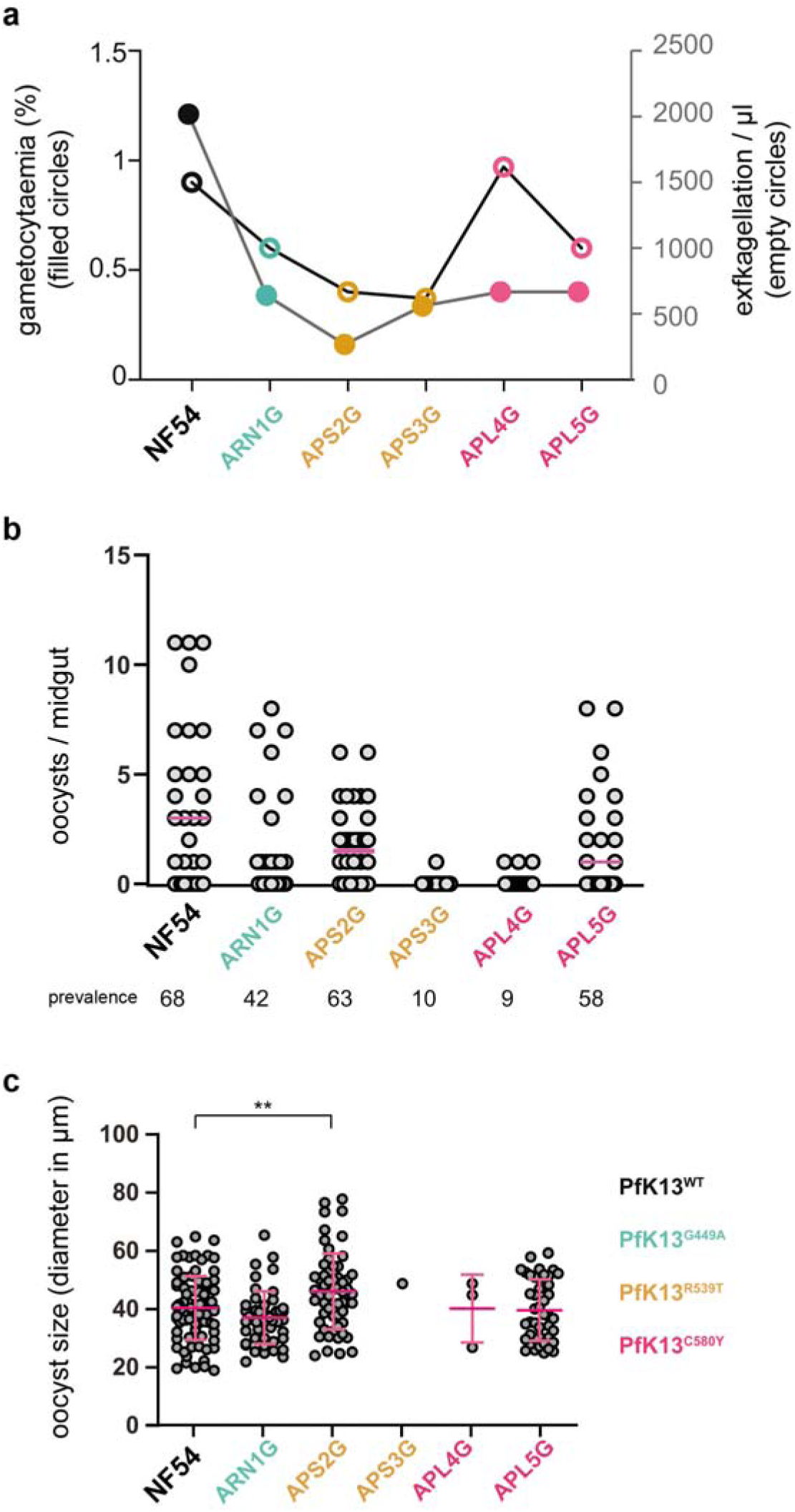
Field isolates form similar-sized oocysts. **A**. Gametocytaemia (represented by filled circles) and exflagellation rates (represented by empty circles) of five field isolates. **B.** *An. stephensi* mosquitoes were infected with field isolates, and oocyst numbers and prevalence are shown. **C.** The size of day 10 oocysts was measured in (**B**). Graph shows diameter of each oocyst found. PfK13 genotype is not related to oocyst size. Isolate APS2G shows significantly bigger oocysts than NF54 (unpaired t-test, ** p<0.01).

**Figure S3.**
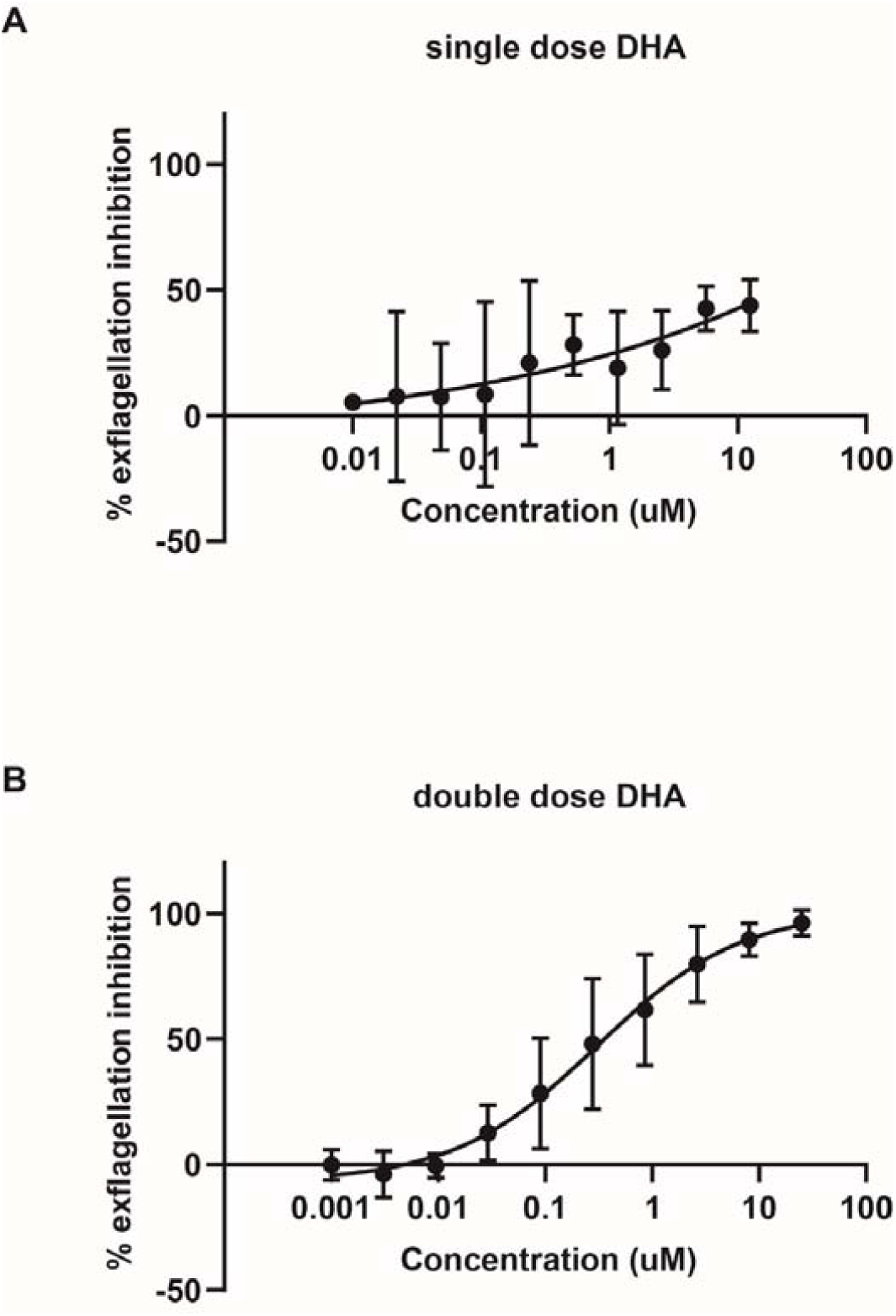
Double-dose DHA elicits a more stable dose-response curve measuring exflagellation inhibition. **A.** NF54 stage V gametocytes were incubated with increasing concentrations of DHA for 24 hours. After incubation, exflagellation centres were counted to give an estimate of % exflagellation inhibition of DHA. Results showed to be very unstable. Three independent biological replicates are shown. **B.** NF54 stage V gametocytes were incubated with increasing concentrations of DHA for 24 hours, after which incubation with the same concentration was repeated for another 24 hours. After 48 hours, exflagellation centres were counted to give an estimate of % exflagellation inhibition of DHA. Results are more stable than with the single dose regimen. Three independent biological replicates are shown

**Figure S4.**
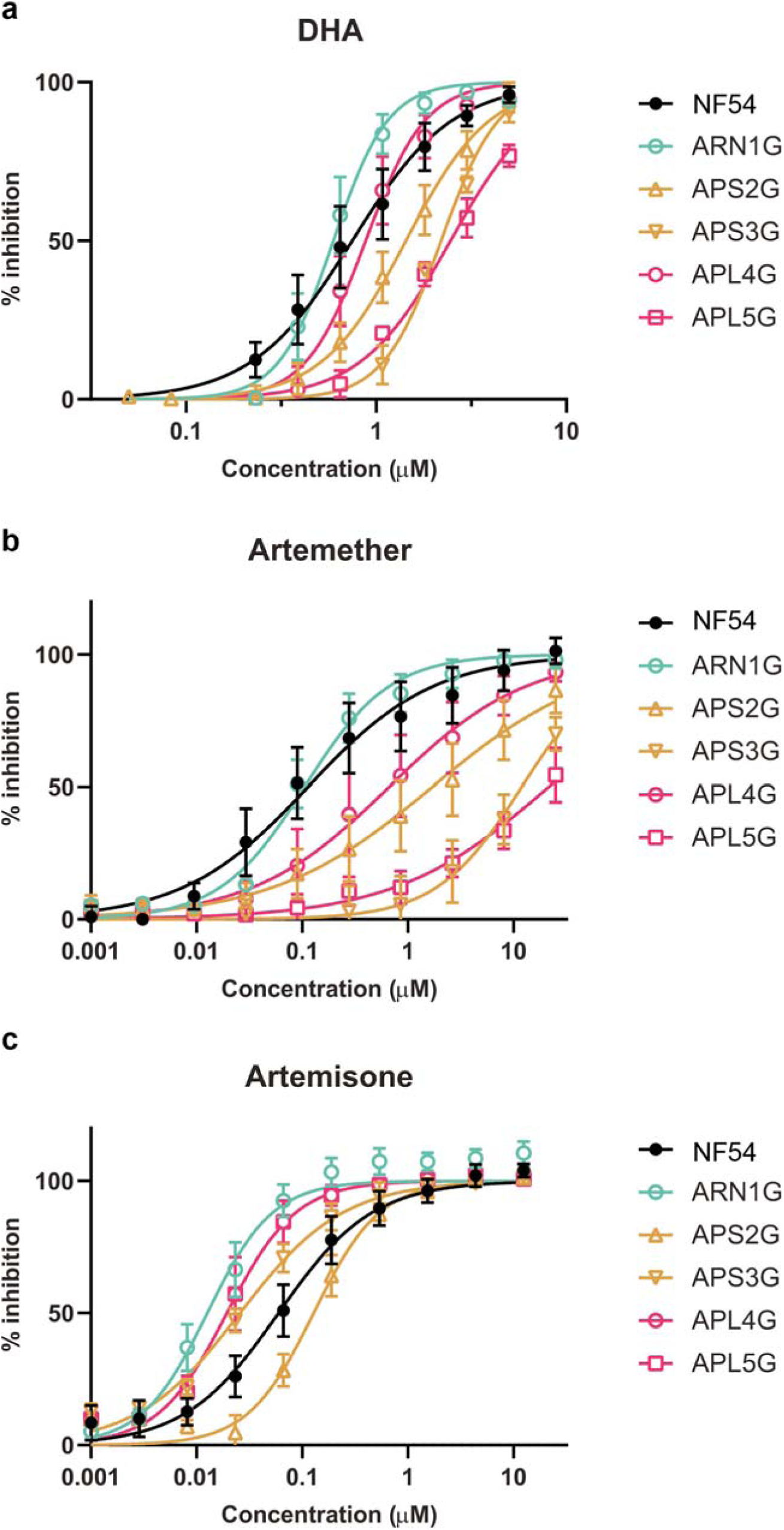
Dose-response curves of three artemisinin-derivatives and their effect on exflagellation inhibition. **A.** DHA. **B.** Artemether. **C.** Artemisone.

**Figure S5.**
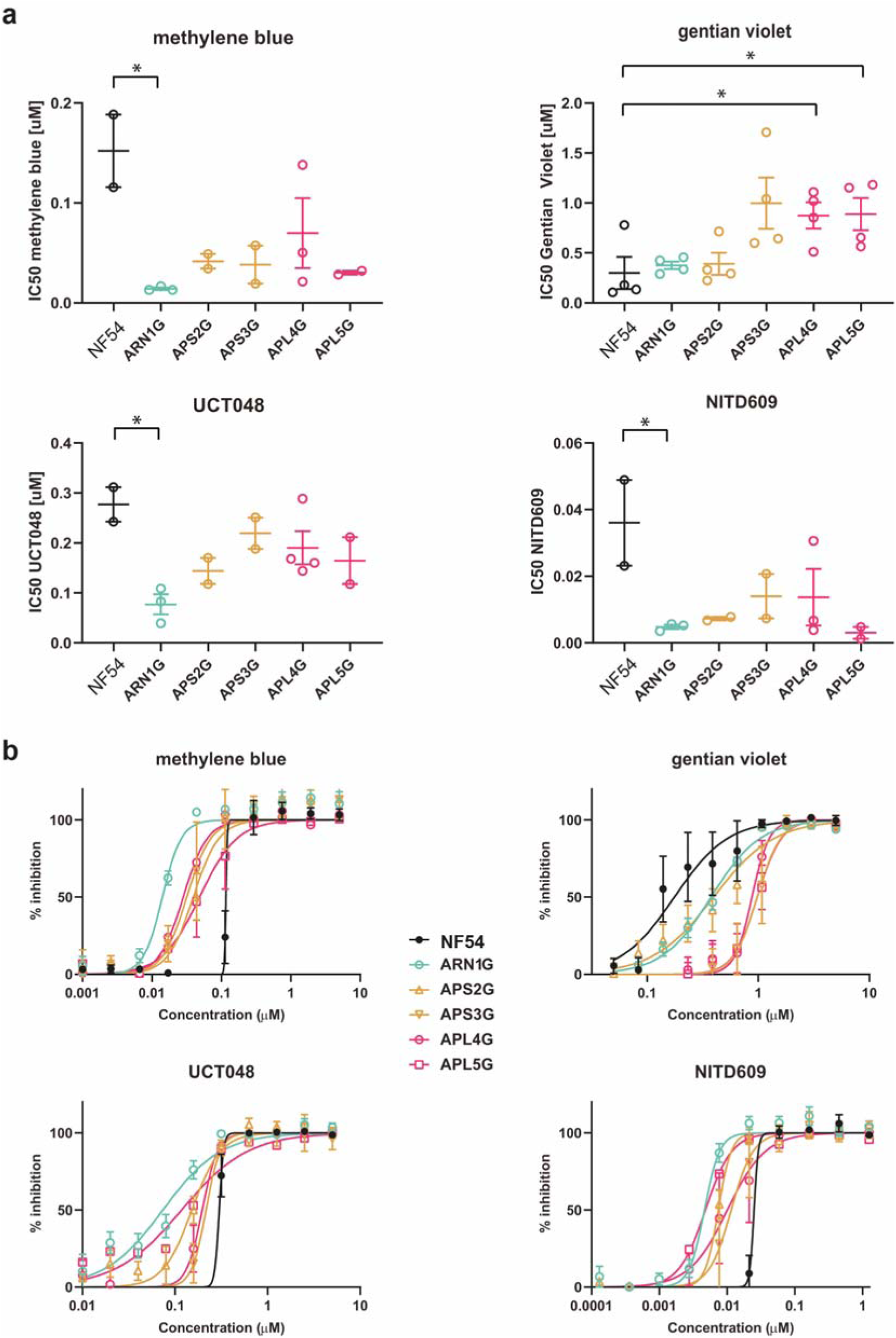
Dose-response curves of four antimalarial compounds and their effect on exflagellation inhibition. **A.** IC_50_ values. **B.** Drug curves.

**Figure S6.**
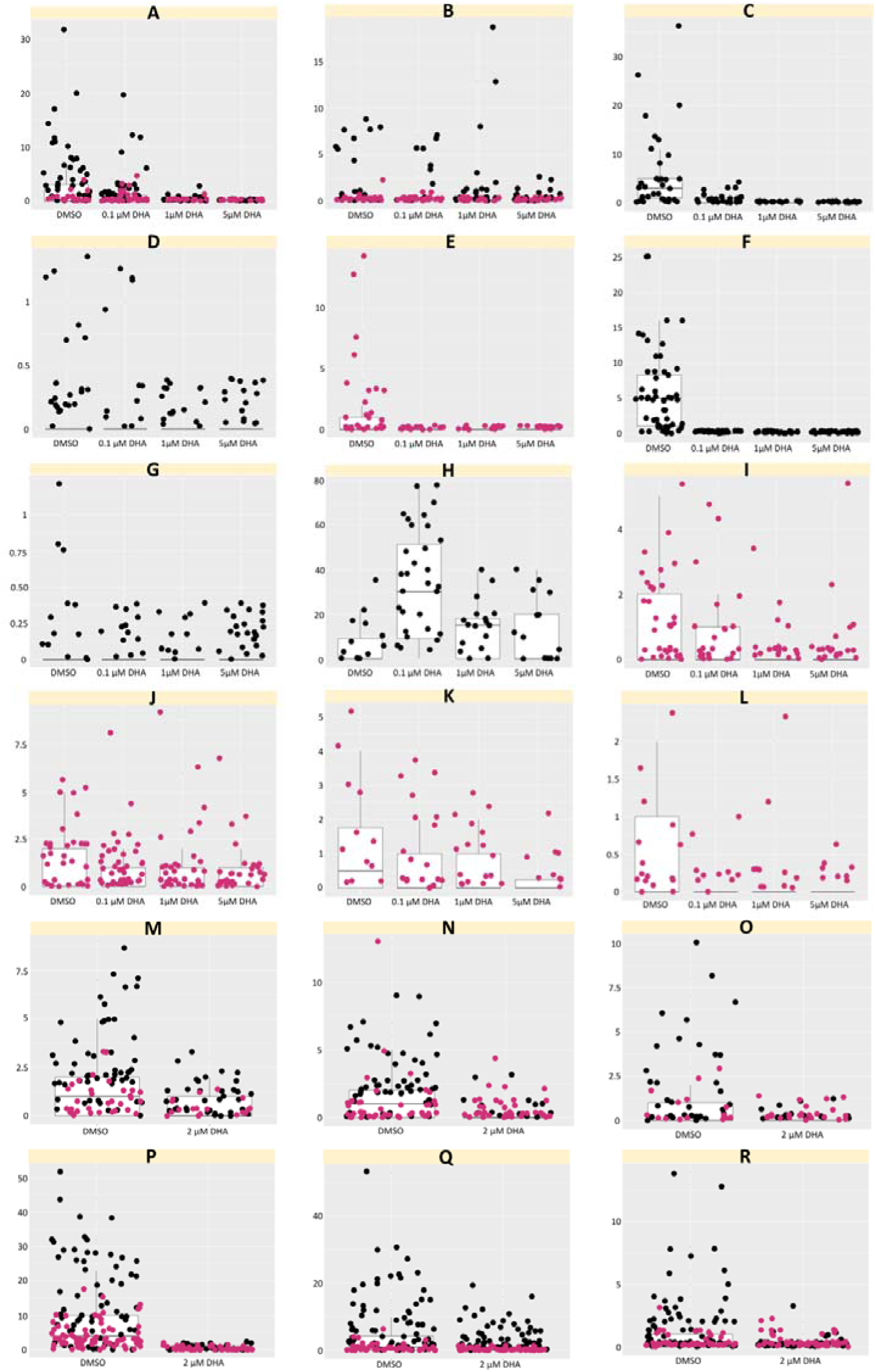
Overall *P. falciparum* infection intensity in 18 individual SMFA (Feed A-R). Each dot represents a single midgut dissected with the number of oocysts plotted on the graph. **A-L.** DMSO, 0.1 µM DHA, 1 µM DHA, and 5 µM DHA were added to **NF54** (blue) and **APL5G** (red). **M-R. I**ncubation for 48 hours pre-feed with DMSO and 2 µM DHA.

## Supplementary Data Sheets (separate files)

Excel spreadsheets of:

1. DGFA data – male exflagellation data
2. SMFA data – oocyst counts

